## Supplementary Table 1, Supplementary Figures 1-11 and the Figure Legends for "Membrane Lipid Nanodomains Modulate HCN Pacemaker Channels in Nociceptor DRG Neurons"

This file includes:

Supplementary Table 1
Supplementary Figures 1-11 and the Figure Legends

Supplementary Table 1

*Gating parameters of native HCN currents in nociceptor DRG neurons (data shown as mean ± s.e.m.)*

| Conditions | 2-sec hyperpolarization protocol | | | 5-sec hyperpolarization protocol | | |
|---|---|---|---|---|---|---|
| | $V_{1/2}$ (mV) | $V_s$ (mV) | ΔG (kcal/mol) | $V_{1/2}$ (mV) | $V_s$ (mV) | ΔG (kcal/mol) |
| control | -92.7 ± 2.5 | 9.1 ± 0.2 | -6.0 ± 0.2 | -85.8 ± 2.3 | 6.6 ± 0.2 | -7.7 ± 0.4 |
| β-CD | -91.3 ± 1.2 | 10.3 ± 0.2 | -5.3 ± 0.1 | -83.2 ± 2.1 | 9.7 ± 0.2 | -5.1 ± 0.2 |
| Paclitaxel | -88.6 ± 1.7 | 10.8 ± 0.5 | -4.9 ± 0.2 | -82.0 ± 4.2 | 8.4 ± 0.4 | -5.8 ± 0.1 |
| Water-Soluble Cholesterol (WSC) | -99.7 ± 1.9 | 9.6 ± 0.2 | -6.1 ± 0.1 | -93.4 ± 2.1 | 7.6 ± 0.3 | -7.3 ± 0.2 |
| +cAMP | -83.3 ± 1.3 | 8.7 ± 0.1 | -5.6 ± 0.1 | -78.5 ± 0.8 | 5.8 ± 0.2 | -8.0 ± 0.3 |
| β-CD + cAMP | -81.2 ± 1.7 | 10.3 ± 0.6 | -4.7 ± 0.2 | -76.0 ± 1.6 | 7.7 ± 0.1 | -5.8 ± 0.3 |
| Paclitaxel +cAMP | -80.2 ± 1.7 | 10.3 ± 0.4 | -4.6 ± 0.1 | -72.9 ± 1.1 | 9.0 ± 0.7 | -4.9 ± 0.4 |
| α-CD | -92.1 ± 4.2 | 9.4 ± 0.1 | -5.8 ± 0.2 | | | |
| SNI-contralateral | -94.0 ± 1.1 | 9.3 ± 0.2 | -6.0 ± 0.2 | | | |
| SNI-ipsilateral | -88.4 ± 1.3 | 10.5 ± 0.2 | -5.0 ± 0.1 | | | |
| SNI-ipsilateral + WSC | -86.7 ± 1.7 | 9.4 ± 0.3 | -5.5 ± 0.2 | N.A. | N.A. | N.A. |
| SNI-contralateral + cAMP | -78.5 ± 2.1 | 7.3 ± 0.3 | -6.4 ± 0.3 | | | |
| SNI-ipsilateral + cAMP | -79.8 ± 3.1 | 9.8 ± 0.3 | -4.8 ± 0.2 | | | |

Figure S1

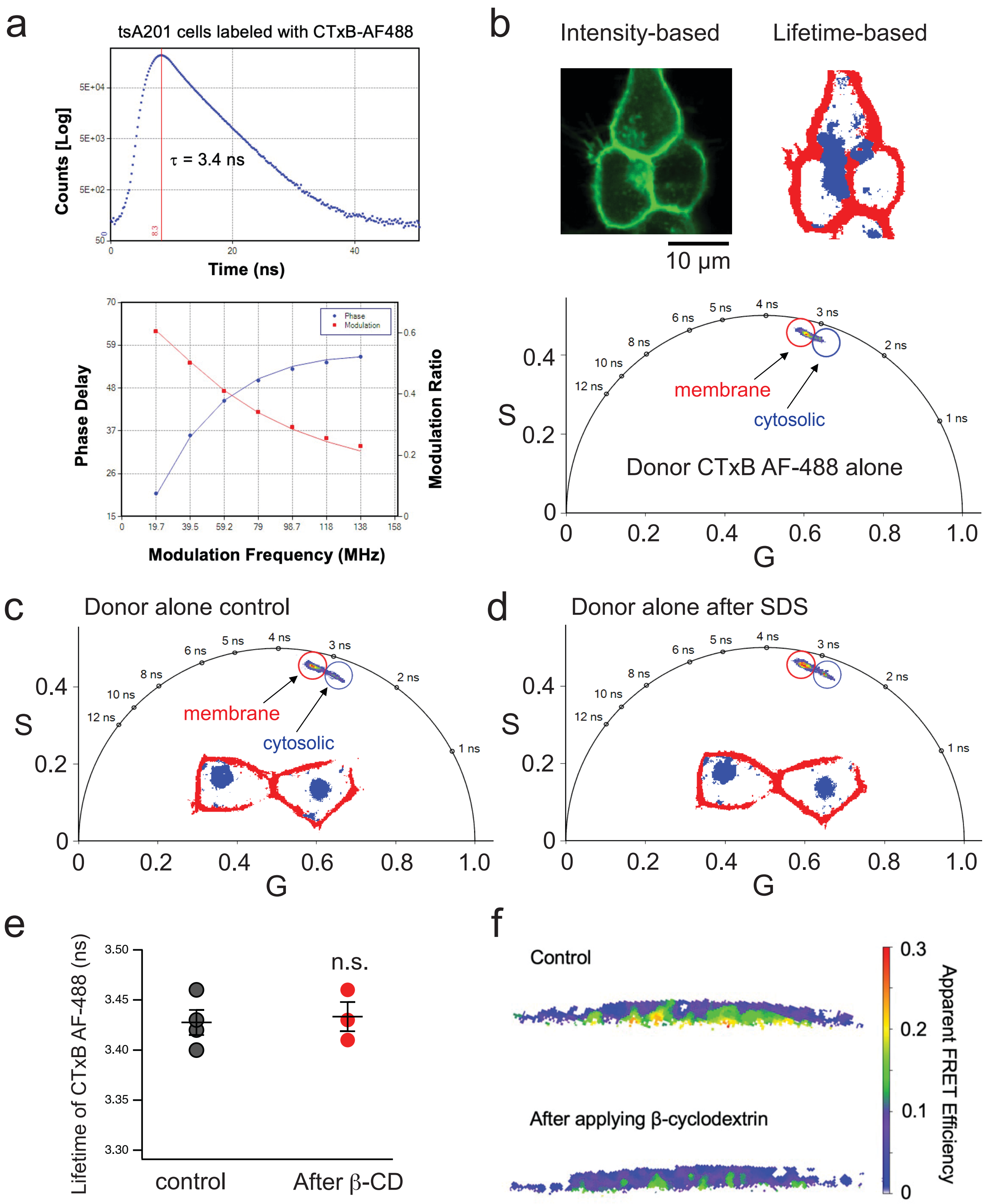

a
tsA201 cells labeled with CTxB-AF488
Counts [Log]
τ = 3.4 ns
Time (ns)
Phase Delay
Modulation Ratio
Phase
Modulation
Modulation Frequency (MHz)
b
Intensity-based
Lifetime-based
10 μm
membrane
cytosolic
Donor CTxB AF-488 alone
S
G
c
Donor alone control
membrane
cytosolic
d
Donor alone after SDS
e
Lifetime of CTxB AF-488 (ns)
n.s.
control
After β-CD
f
Control
After applying β-cyclodextrin
Apparent FRET Efficiency

**Supplementary Figure 1. Frequency-domain phasor FLIM approach for imaging OMDs in living cells.**

(**a**) Time-domain and frequency-domain measurements of FLIM imaging of cholera toxin subunit B (CTxB) Alexa Fluor 488 (AF-488) conjugate (20 nM pentamers applied to tsA201 cells). The modulation ratio decreases with an increase of the modulation frequency of the excitation laser while the phase delay increases with it. (**b**) Representative intensity-based and lifetime-based images of a cluster of cells labeled with the FRET donor CTxB AF-488 alone. Cytosolic (blue) CTxB probes that are internalized can be distinguished from the membrane-localized (red) CTxB in their lifetime displayed on the phasor plot. (**c**) Exemplar phasor plot of CTxB AF-488 showing the separation of lifetimes of the membrane (red) and cytosolic (blue) species in dPBS. Two cells with relatively high amounts of internalized CTxB were selected to highlight the characteristics of the phasor plot in dissecting different lifetime species. **(d)** Lifetimes of membrane (red) and cytosolic (blue) CTxB AF-488 probes after acute application of SDS. SDS did not change the lifetime of membrane-localized CTxB AF-488. (**e**) Summary data of CTxB AF-488 lifetime before and after application of β-cyclodextrin. β-cyclodextrin treatment did not change the lifetime of the CTxB AF-488 at the plasma membrane; $n = 3 – 4$, $p = 0.77$. (**F**) Heatmap images of the localized FLIM-FRET efficiency of the plasma membrane of Xenopus oocytes. β-cyclodextrin decreased the FRET efficiency.

Figure S2

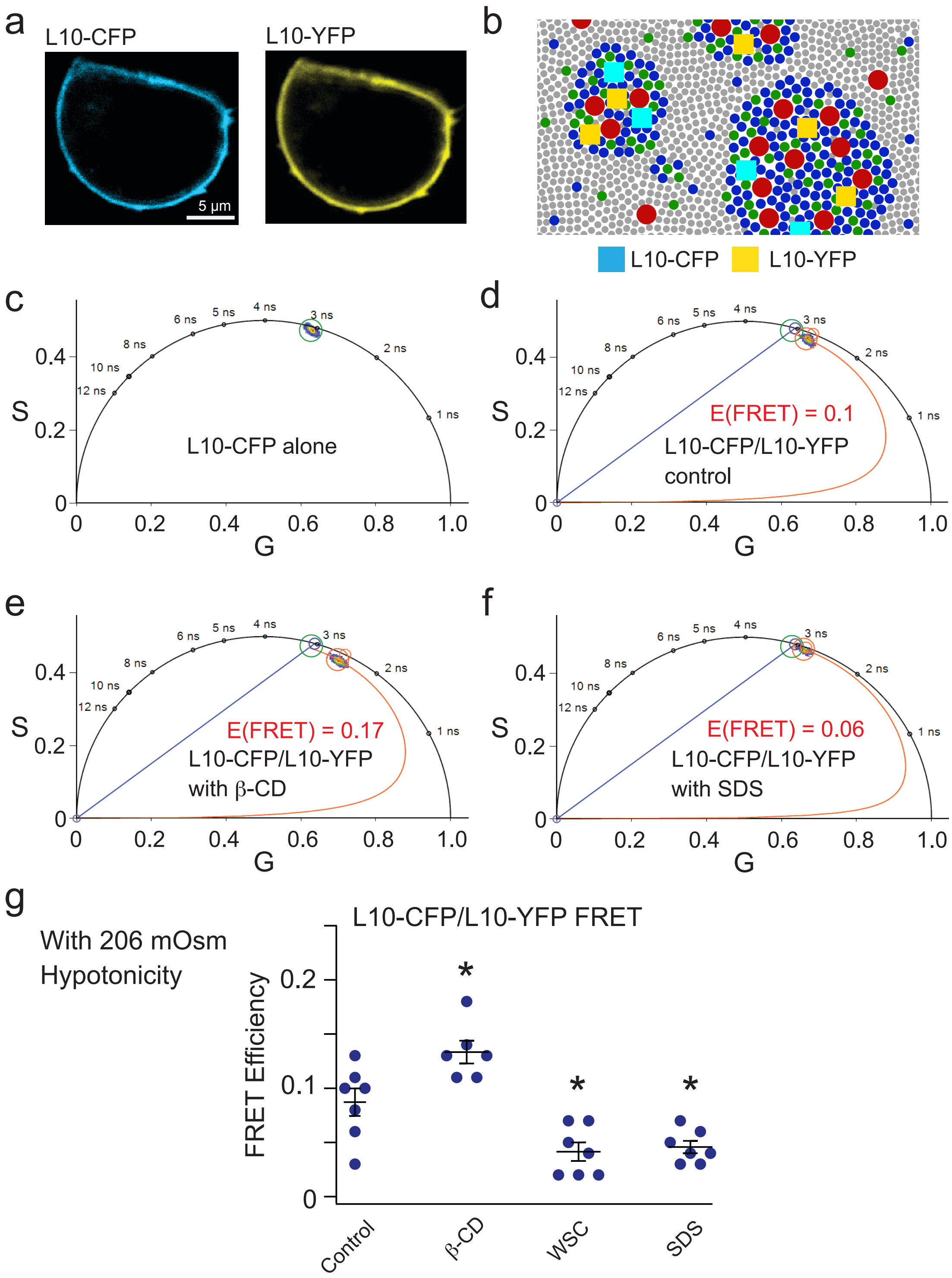


**Supplementary Figure 2. FLIM-FRET between OMD probes L10-CFP and L10-YFP.**

(**a**) Representative fluorescence images of the L10-CFP and L10-YFP probes in tsA201 cells. (**b**) Illustration of ordered membrane domains, where L10 probes are represented by cyan (CFP) and yellow

(YFP) squares. (**c**) Representative phasor plot of fluorescence lifetime of L10-CFP donor alone. (**d-f**) Representative phasor plots of FRET between the L10-CFP probe (donor) and L10-YFP probe (acceptor) at the control condition (d), after acute application of 5 mM β-CD (e), and after acute application of 50 μM SDS (f). (**g**) Summary data of the same L10-CFP/L10-YFP FLIM-FRET experiments applying respective treatments, with 206 mOsm hypotonic pre-treatment. Data shown are mean ± s.e.m., n = 6 – 7, *p = 0.015 for the β-CD condition, *p = 0.012 for the WSC condition, and *p = 0.025 for the SDS condition.

Figure S3

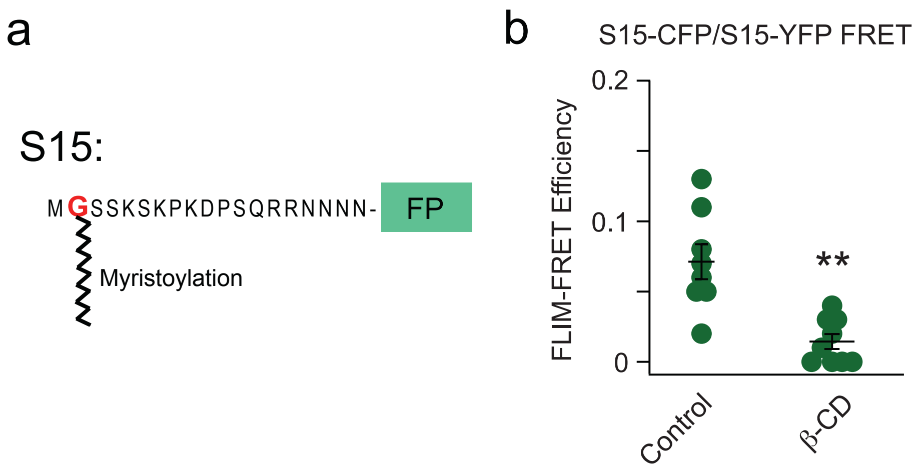


**Supplementary Figure 3. FLIM-FRET between the fluorescent membrane probes S15-CFP and S15-YFP.**

(**a**) Cartoon showing the S15 probe, with only one myristylation site that facilitates the localization of S15 to disordered membrane regions. (**b**) Summary of the FRET efficiency between S15-CFP and S15-YFP with and without β-CD. Data shown are mean ± s.e.m., n = 8 for the control and n = 9 with β-CD, **p = 5e-4.

Figure S4

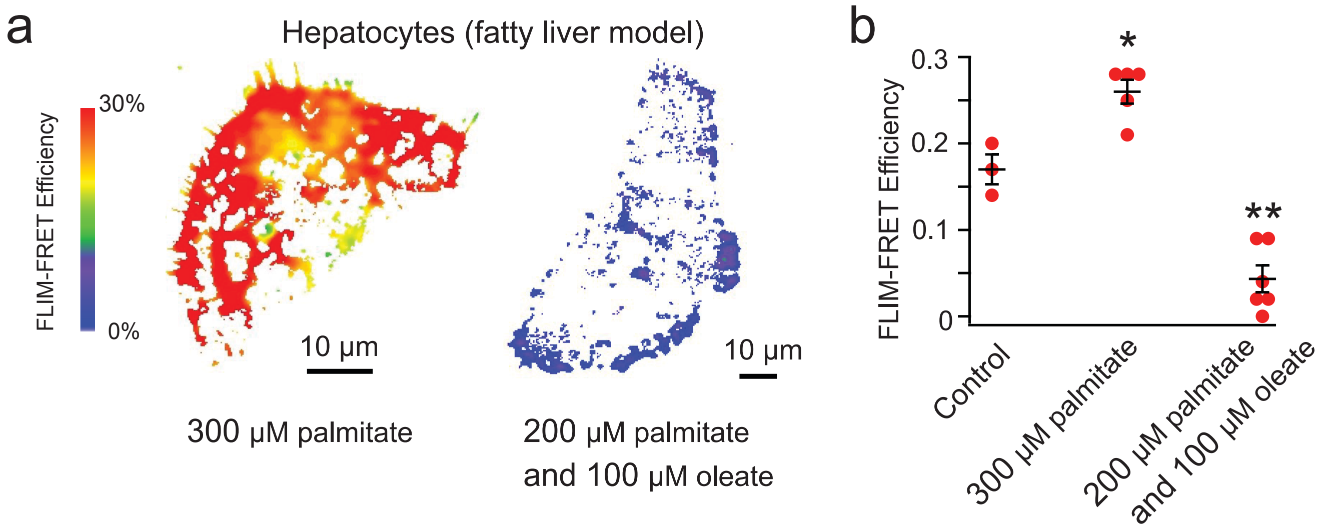


**Supplementary Figure 4. Lipid tail structure affects the OMD size in hepatocytes.**

(**a**) Representative heatmap images of mouse hepatocytes with different fatty acid treatments. (**b**) Summary data of the effects produced by the palmitate supplementation versus the combination of palmitate and oleate supplementations as shown in panel a. Data shown are mean ± s.e.m., n = 3 - 6, *p = 0.011 with palmitate and **p = 8e-4 with both palmitate and oleate compared to the control.

Figure S5

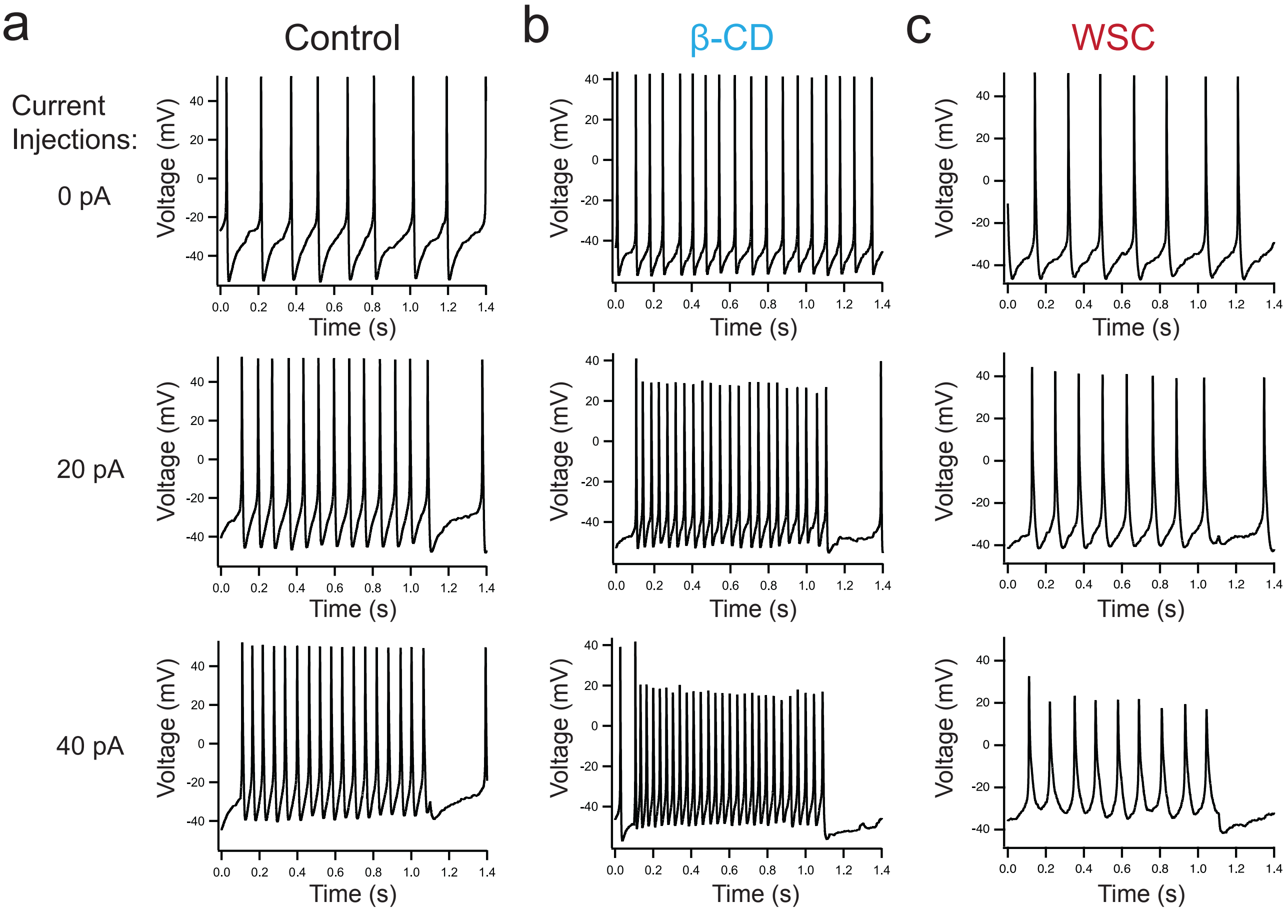


**Supplementary Figure 5. Action potential firing elicited by current injections for small DRG neurons.**

(**a-c**) Representative spontaneous action-potential firings of the control nociceptor DRG neurons (a) and those treated by β-cyclodextrin (b) and WSC (c), with different amplitudes of current injections.

Figure S6

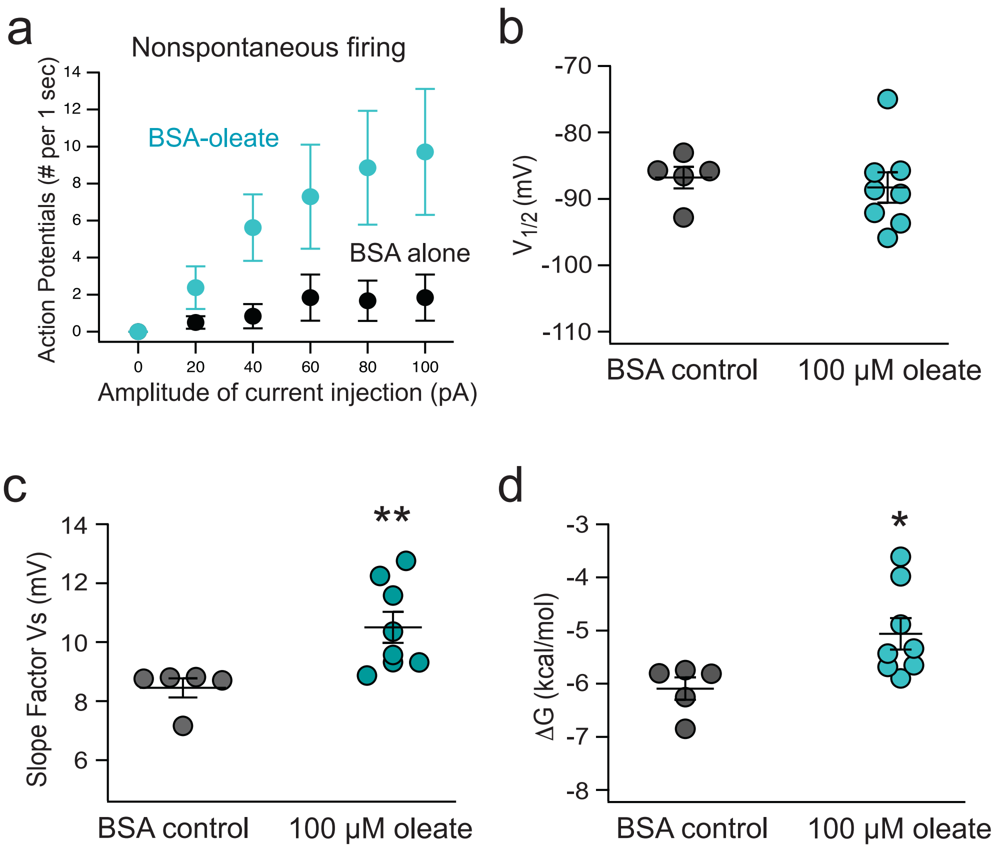


**Supplementary Figure 6. Summary of the effects of the oleate treatment on the action potential firing and gating parameters of native HCN channels in small DRG neurons.**

**(a)** Summary data of the effect of 100 μM oleate supplementation on the action potential firing in response to injected currents for nonspontaneous small DRG neurons. Data are presented as mean ± s.e.m. (n = 6 - 8). Statistical significance is reported as follows: p = 0.197 (+20 pA), *p = 0.047 (+40 pA), p = 0.123 (+60 pA), p = 0.06 (+80 pA), and p = 0.07 (+100 pA) for the respective current injections. (**b-d**) Summary data of the effect of oleate supplementation on gating parameters: the $V_{1/2}$ for the G-V relationship (panel b, p = 0.65), the slope factor (panel c, **p = 0.002), and the free energy required for channel activation (panel d, *p = 0.03) of HCN2 channels. Data are shown as mean ± s.e.m., n = 5 for the BSA control, and n = 8 with oleate.

Figure S7

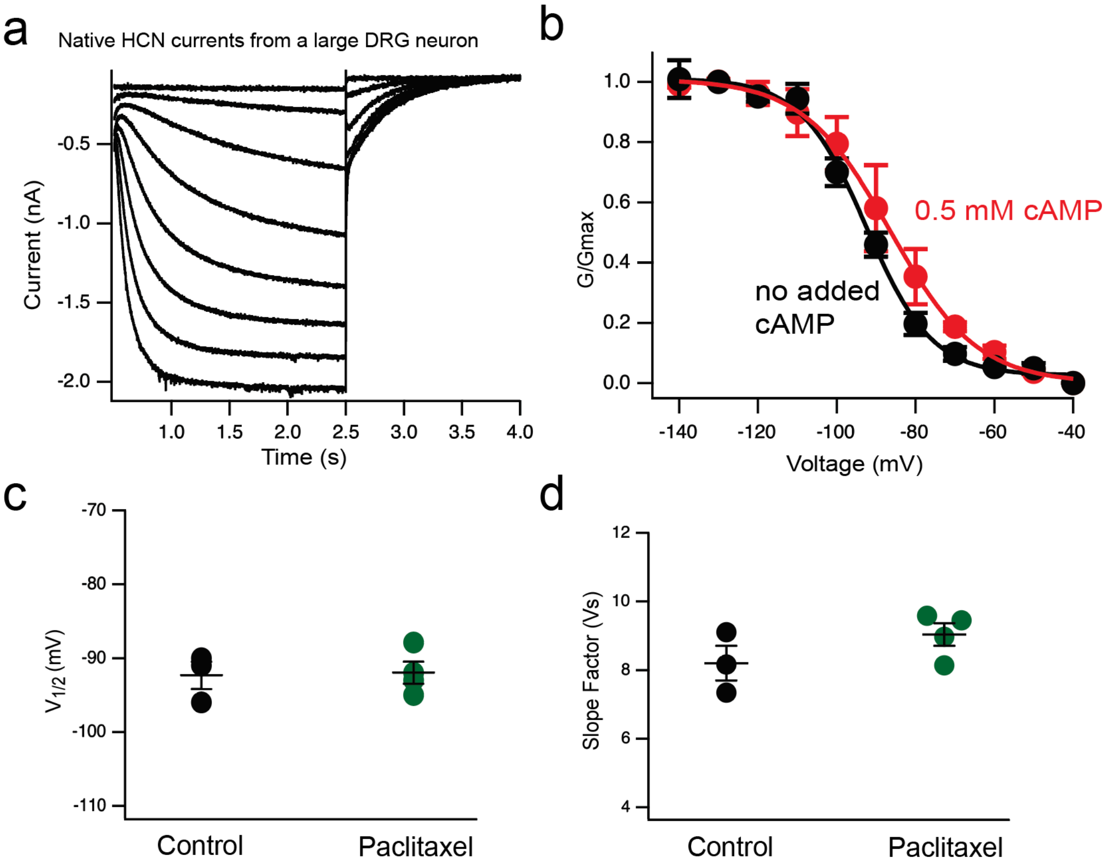


**Supplementary Figure 7. Effects of the paclitaxel treatment on the gating parameters of native HCN channels in large DRG neurons.**

(**a**) Representative endogenous HCN currents of large DRG neurons elicited by a series of hyperpolarizing voltage pulses. (**b**) Conductance-voltage relationships of the endogenous HCN currents in large DRG neurons with (red) or without (black) the added 0.5 mM cAMP in the pipette solution. n = 3 - 4. The averaged $V_{1/2}$ values are -92.3 ± 1.8 mV without added cAMP (n = 3) and -87 ± 4.6 mV with the 0.5 mM cAMP, n = 4, p = 0.28. (**c-d**) Summary data of the effect of 10 nM paclitaxel treatment on the gating parameters $V_{1/2}$ (c) and the slope factor $V_s$ (d) of endogenous HCN currents of large DRG neurons. Data are shown as mean ± s.e.m., p = 0.88 for the $V_{1/2}$, p = 0.2 for the slope factor.

Figure S8

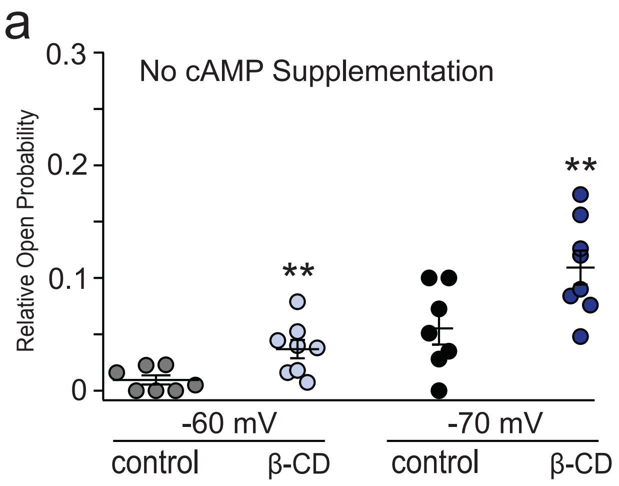


**Supplementary Figure 8. Summary of the effects of β-cyclodextrin treatment on the normalized open probability of native HCN channels in small DRG neurons.**

(**a-b**) Summary data of relative open probability based on the normalized G-V relationships in the presence and absence of β-CD at -60 mV and -70 mV with no added cAMP (a) and 0.5 mM cAMP (b). Data are shown as mean ± s.e.m., n = 5 - 8, **p < 0.01 compared to the control for each condition, p = 0.17, no statistical significance (n.s.).

Figure S9

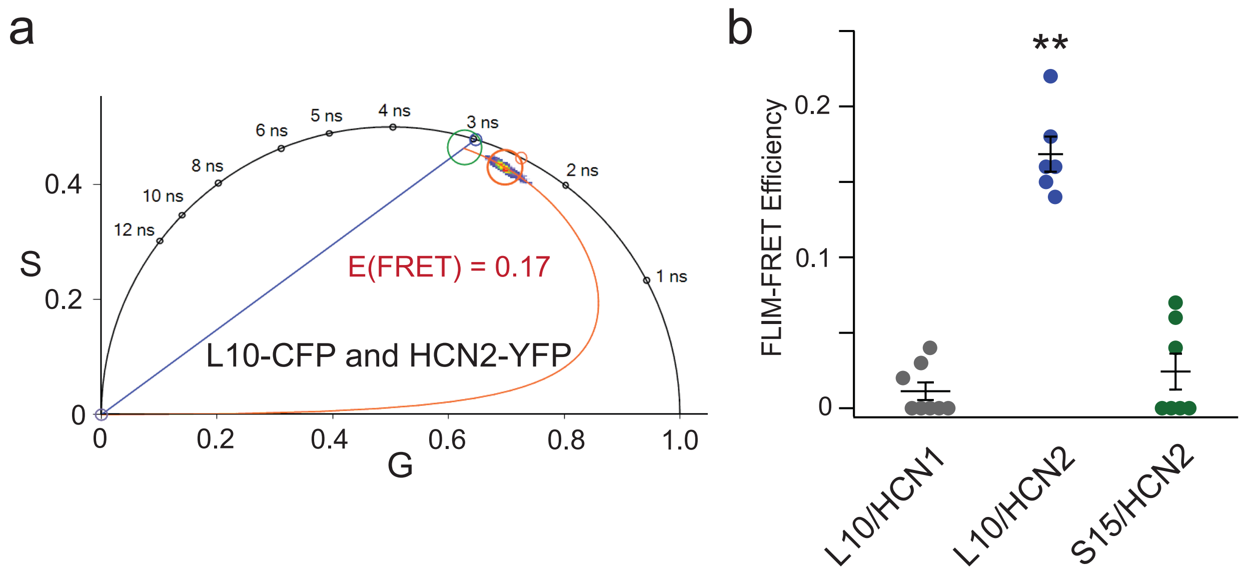


**Supplementary Figure 9. Phasor FLIM-FRET between the OMD probe L10-CFP, S15-CFP, and hHCN2-YFP.**

(**a**) Representative phasor plots of FRET between L10-CFP with hHCN2-YFP. The green cursor indicates the lifetime of cells transfected with L10-CFP alone. (**b**) Summary data of the FRET efficiency involving L10-CFP with hHCN1-YFP and hHCN2-YFP, and S15-CFP with hHCN2-YFP. HCN1-YFP, serving as a negative control, exhibits a diminished propensity for localization within the OMDs. Data are shown as mean ± s.e.m., n = 6 – 8, $^{*}p < 0.05$; $^{**}p < 0.01$.

Figure S10

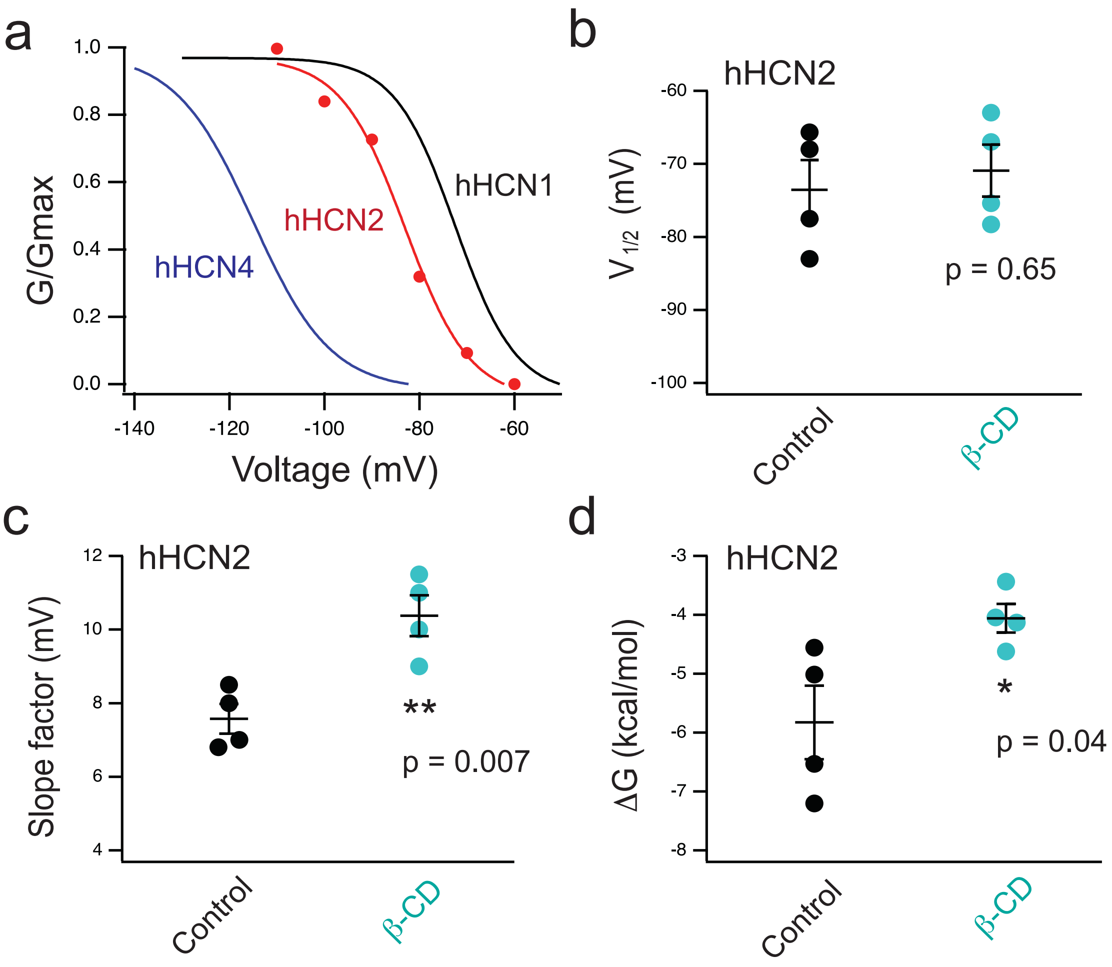


**Supplementary Figure 10. Gating properties of human HCN2 channels and their regulation by β-CD, as measured in tsA-201 cells.**

(**a**) Representative G-V relationships of human HCN2 (red) compared to human HCN1 (black) and HCN4 (blue) channels. The HCN1 and HCN4 data were adapted from previous research from our lab. The $V_{1/2}$ = -115 mV (hHCN4), -83 mV (hHCN2), and -72.5 mV (hHCN1); The slope factor Vs = 8.2 mV (hHCN4), 6.8 mV (hHCN2), 6.4 mV (hHCN1); the free energy for activation $\Delta G$ = -8.3 kcal/mol (hHCN4), -7.2 kcal/mol (hHCN2), and -6.7 kcal/mol (hHCN1). (**b-d**) Summary data of the effect of 5 mM β-CD application on gating parameters for hHCN2 in tsA cells: the $V_{1/2}$ for the G-V relationship (b), the slope factor (c), and the free energy required for channel activation (d) of HCN2 channels. The hHCN2 currents were recorded using the cell-attached mode on tsA cells, with a similar hyperpolarization protocol. Data are shown as mean ± s.e.m., n = 4.

Figure S11

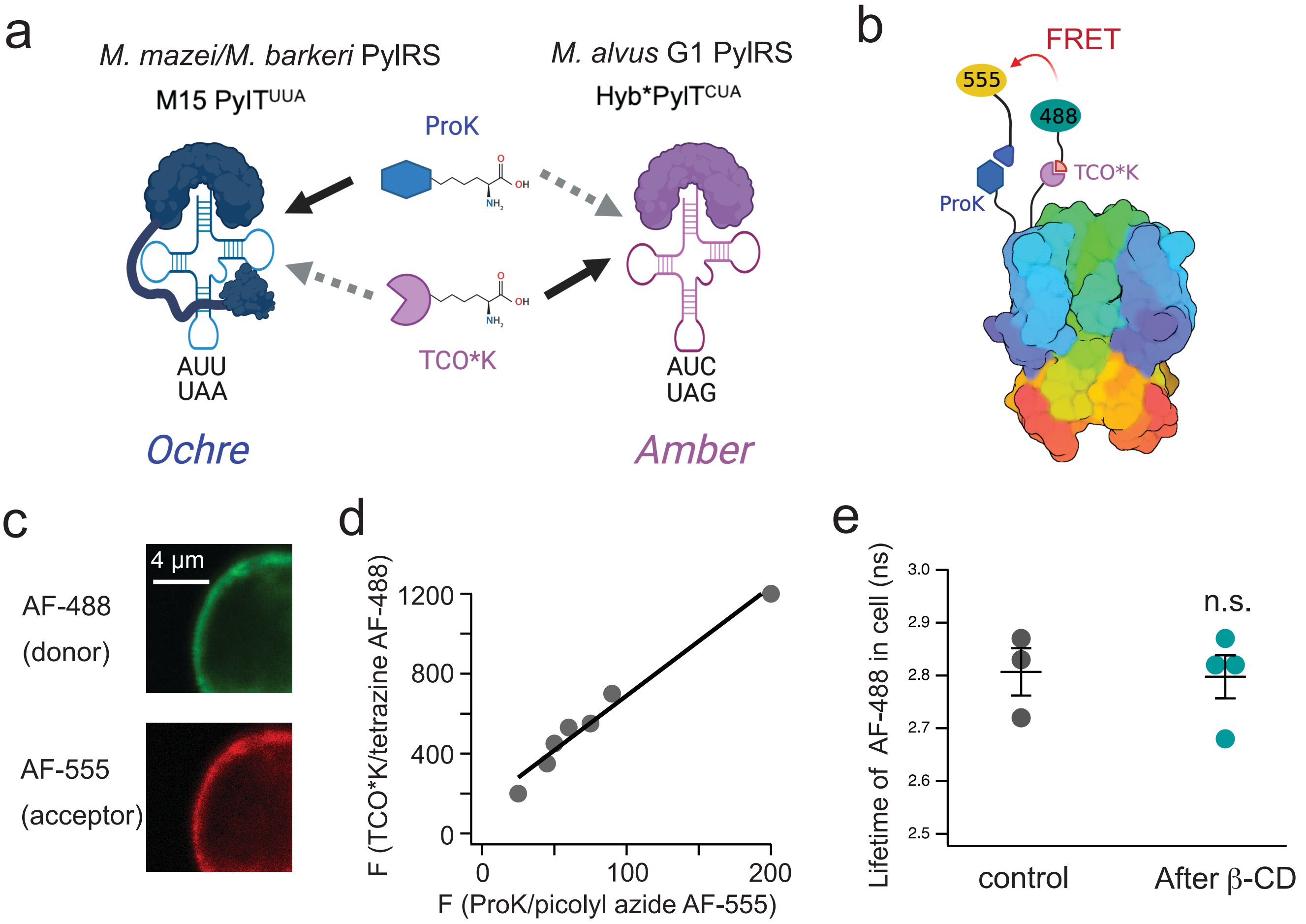


**Supplementary Figure 11. Strategies for incorporating two different noncanonical amino acids to HCN channels orthogonally.**

(**a**) The system incorporated a G1 PylRS-Y125A variant in combination with an enhanced G1 tRNA. The improved tRNA features two mutations (A41AA and C55A) and includes an acceptor stem portion derived from the Pyl tRNA of M. alvus Mx1201, known as hybPylT. Additionally, we used tRNA (M15-PylT)/PylRS pairs from M. mazei/M. barkeri (Mma). It is noteworthy that when applied in mammalian cells, the tRNA/PylRS from M. alvus does not exhibit cross-reactivity with the tRNA/PylRS from M. mazei/M. barkeri. The hybPylT tRNA facilitates amber stop-codon suppression, encoding TCO*K, while the M15-PylT tRNA facilitates ochre stop-codon suppression, encoding N-propargyl-L-lysine (ProK). (**b**) Cartoon illustrating the labeling of HCN channels with the dual stop-codon suppression system and click chemistry reactions CuAAC and IEDDA. (**c**) Representative images of a tsA201 cell expressed with the hHCN2 L323TAG/T240TAA construct that is dually labeled with AF-488 and AF-555. (**d**) Linear correlation of the labeled AF-488 versus AF-555 fluorescence using the hHCN2 L323TAG/T240TAA construct. (**e**) Summary data of TCO*K linked tetrazine AF-488 lifetime at the plasma membrane before and after application of β-cyclodextrin. β-cyclodextrin treatment did not change the lifetime of the AF-488. Data are shown as mean ± s.e.m., n = 3 – 4, p = 0.89, no statistical significance (n.s.).
